# Visually-guided compensation of deafening-induced song deterioration

**DOI:** 10.1101/2024.11.04.621802

**Authors:** M Rolland, A Zai, RHR Hahnloser, C Del Negro, N Giret

**Author notes:** Equal contributions (M Rolland, A Zai).

## Abstract

Human language learning and maintenance depend primarily on auditory feedback but are also shaped by other sensory modalities. Individuals who become deaf after learning to speak (post-lingual deafness) experience a gradual decline in their language abilities. A similar process occurs in songbirds, where deafness leads to progressive song deterioration. However, songbirds can modify their songs using non-auditory cues, challenging the prevailing assumption that auditory feedback is essential for vocal control. In this study, we investigated whether deafened birds could use visual cues to prevent or limit song deterioration. We developed a new metric for assessing syllable deterioration called the spectrogram divergence score. We then trained deafened birds in a behavioral task where the spectrogram divergence score of a target syllable was computed in real-time, triggering a contingent visual stimulus based on the score. Birds exposed to the contingent visual stimulus—a brief light extinction—showed more stable song syllables than birds that received either no light extinction or randomly triggered light extinction. Notably, this effect was specific to the targeted syllable and did not influence other syllables. This study demonstrates that deafness-induced song deterioration in birds can be partially mitigated with visual cues.

## Introduction

Human language and speech learning rely on vocal learning processes that involve imitation and sensory-motor integration. Infants learn to speak by imitating the vocal communication signals of the individuals around them. During development, an individual’s vocalizations become more and more accurate, guided by the sensory, *i.e*. auditory, feedback of her own voice. Infants born deaf or with severe hearing impairments never acquire typical adult speech. In adults, post-lingual deafness — the complete loss of hearing abilities after acquisition of speech sounds — leads to a degradation of speech, including the loss of phonetic precision of vowels and consonants and the change in duration of syllables and sentences (Cowie et al., 1982; Leder et al., 1987c, 1987a, 1987b; Waldstein, 1990; Lane and Webster, 1991). Auditory feedback thus plays a critical role for adult speech maintenance. Sharing close similarities with speech learning in humans, learning and maintenance of the vocal signals produced by songbirds also critically rely on the processing of auditory feedback. Deaf juvenile songbird never manage to develop typical song of their own species (Konishi, 1964, 1965a, 1965b, 2004; Iyengar and Bottjer, 2002). Intact hearing abilities are thus required for a juvenile to learn to imitate the songs of adult conspecifics. Once adult, songbirds keep on processing the auditory feedback that allows them to adjust their vocal behaviour when facing environmental noise constraints (Okanoya and Yamaguchi, 1997; Leonardo and Konishi, 1999; Brainard and Doupe, 2000; Tumer and Brainard, 2007; Andalman and Fee, 2009; Sober and Brainard, 2009; Derryberry et al., 2020). Deafened adult songbirds exhibit a progressive and dramatic degradation of their vocal production (Nordeen and Nordeen, 1992, 2010; Woolley and Rubel, 1997; Lombardino and Nottebohm, 2000; Horita et al., 2008; Wittenbach et al., 2015).

Auditory feedback is thus critical for song maintenance, even in closed-ended learner species such as zebra finches (*Taeniopygia guttata*) who produce highly stereotyped song syllables once adults (>90 days post-hatch, dph). Interestingly however, adult male zebra finches are able to adjust a song syllable using non-auditory feedback signals, *i.e*. a visual signal such as a transient light extinction (Zai et al., 2020) or mild subcutaneous electric stimulations (McGregor et al., 2022). The behavioural training paradigm relies on the contingent delivery of the sensory signal depending on the pitch of a song syllable. Importantly, deaf birds modified the pitch of the selected song syllable in order to get more transient light extinction while hearing birds did the opposite (Zai et al., 2020). Furthermore, the magnitude of pitch change was much higher in deaf than hearing birds. The intrinsic value of the transient light extinction could therefore be assumed to differ between deaf and hearing birds, being attractive for deaf birds while repulsive for hearing birds, respectively

As deaf birds adapt the pitch of a selected song syllable using information provided by a visual signal (Zai et al., 2020), we wondered whether a similar behavioural protocol could be used to prevent or at least delay deafening-induced song degradation. Because deafening-induced song degradation includes a wide range of spectro-temporal changes in the structure of song syllables (Nordeen and Nordeen, 1992; Wang et al., 1999; Brainard and Doupe, 2001; Horita et al., 2008), syllables uniformly and gradually get noisier (Horita et al., 2008; Hamaguchi et al., 2014), — we aimed to examine whether deafened birds could use non auditory signals to control the acoustic structure of a selected song syllables. To do so, we first developed a metric for measuring syllable acoustic stability and for estimating in real time the degree of vocal deterioration. We then modified the operant conditioning paradigm based on the pitch-contingent delivery of a transient light extinction (Zai et al., 2020) such that the visual signal was delivered depending on the measure of the syllable acoustic stability.

## Method

### 1. Subject and groups

We used 40 adult male (>90 days post-hatch) and young adult (70 days post-hatch) male zebra finches (*Taeniopygia guttata*) raised in our breeding colonies in Orsay and Saclay (France) or in Zurich (Switzerland). All experiments have been approved by the French Ministry of Research and ethical committee “Paris-Sud et Centre (CEEA n°59, project 2017-12) or by the Cantonal Veterinary Office of the Canton of Zurich, Switzerland (license numbers 207/2013 and ZH077/17) and comply with the EU Council Directive 2010/63 of the European Parliament on the protection of animals used for scientific purposes.

Throughout the experiments, the birds were housed individually in sound-proof chambers under a 14-hour day-10-hour night photoperiod cycle, with access to food and water *ad libitum*. Birds resumed singing at a normal rate after 2–5 days in the experimental environment. Chambers were equipped with a wall-attached microphone. Sound signals were band-pass filtered, and digitized at a sampling rate of 32 kHz and songs were detected online using the RecOOrder software (Herbst et al., 2023) developed on LabView (National Instruments, Inc).

Birds were divided in four groups. They included one group of hearing control birds (n=7) and three groups of deafened birds : deaf LO (n=13), deaf random LO (n=5) and deaf no LO (n=15) birds. Deaf LO birds were exposed to the syllable-contingent delivery of a transient light extinction while deaf random LO birds were exposed to a similar amount of transient light extinction than deaf LO birds, but the visual signal exposure did not depend on what the bird was singing. Deaf no LO and hearing birds were not exposed to transient light extinction during the entire course of the experiment.

### 2. Deafening procedure

All birds except hearing birds underwent the deafening procedure. To do so, we applied a technique consisting in the bilateral cochlear ablation (Schwartzkopff, 1949; Zai et al., 2020). It results in the complete and irreversible suppression of the auditory feedback. The birds were anesthetized by inhalation of a mixture of oxygen and isoflurane (induction: 2-3%; maintenance: 1-2%). Once the flexion reflex was no longer observable, the bird was placed in the stereotaxic apparatus in the prone position and its beak is placed at 90° to the horizontal axis. The feathers are removed between the two ears, at the lower part of the skull at the back of the bird’s head. Disinfectant (Vetedine) and local anesthetic (Lurocaine) were applied to the skin of the skull 10 minutes before incising it. The skin was incised over 5 millimeters at the level of the hyoid bone in the antero-posterior direction, in order to expose the neck muscles. The muscles were gently pushed down the skull to expose the cranial surface where the semicircular canals are visible through the skull. The craniotomy was performed with forceps just below the point where the posterior and external semicircular canals cross. Under a light microscope, the topography of the area was observed to determine the position of the dome forming the upper part of the bony canal containing the cochlea, above the bony crest, at a position anteromedial to the oval window. At this dome, a small window was opened with forceps to allow removal of the cochlea. The cochlea was removed from the cavity with a custom-made tungsten hook. After removal, the cochleae including the lagenas were photographed to verify their integrity. The operation was performed bilaterally to induce total deafness by removing both cochleae.

### 3. Visual substitution task of syllable similarity: LO protocol

We ran a custom-made LabView (National Instruments, Inc) program to provide visual substitution contingent on syllable similarity to deafened birds. We targeted a harmonic syllable, later called “target syllable”, using a two-layer neural network (perceptron) trained on a subset of manually clustered vocalizations recorded. We first computed the spectrogram of each rendition of the target syllable produced on the last baseline day before the start of the LO protocol. We applied a Hamming windowing with a sample size of 512 (32 kHz sampling rate) on the raw sound signal. The windowed signal is then transformed into a linear power sound spectrogram using the fast Fourier transform. This is computed on segments of 512 samples with an overlap of 128 samples (corresponding to 4 ms). The resulting spectrogram has a resolution of 4 ms per column and 63 Hz per row. To compute the reference spectrogram, we selected a window of the last 48 ms before the detection carried by the perceptron. We applied a high-pass filter on the sound signal to keep frequencies above 630 Hz (low frequencies mostly contain non-vocal sounds). For each rendition, we thus obtained a matrix of 12 columns (4 ms/column), and 128 rows (63 Hz/row) covering a frequency bandwidth of 630 to 8694 Hz. From each cell of the matrix, we can infer the time (which column), the frequency (which line) and the sound amplitude (cell value), the latter being normalized to the maximum sound amplitude of the matrix. We computed the average reference spectrogram of the target syllable by computing the mean matrix of all the matrices obtained for each rendition of the target syllable.

After deafening, our custom-made program computed a similarity score between the average reference spectrogram of the target syllable and the spectrogram (computed as before) of each rendition of the target syllable sang by the bird. This score, that we call “Spectrogram divergence score”, corresponded to the Euclidian distance computed on a fixed duration window of a 48 ms (12 columns of 4 ms and 128 rows of 63 Hz) part of the target syllable. The score thus provides a measure of syllable similarity. Twelve milliseconds following syllable similarity measurement, we provided the visual substitution that consisted in a transient light extinction (light-off, LO) with a duration in the range of 100 to 500 ms in the housing chamber of the bird either in contingency to the spectrogram divergence score, *i.e*. when the score was lower than a manually set threshold, for the deaf LO birds or not-contingent to the similarity score for the deaf random LO birds. Deaf control and hearing birds were never exposed to transient light extinction during the day. The task started when a bird was producing at least 400 song motifs per day on 3 consecutive days post-deafening. Birds were then involved in the task during 6 consecutive weeks.

### 4. Song analyses

Song data were processed offline in a custom program written with Matlab (v2023b). We performed a manual clean-up of the cluster containing the automatically detected part of the target syllables (window of 48 ms) in order to remove any false positives from the dataset of each recording day. We then computed the spectrogram of each rendition of the target syllable part and applied a normalization to the maximum amplitude for each spectrogram in order to compensate for any environmental variability (e.g. bird position toward the microphone, background noise, *etc*.). We then computed the spectrogram divergence score as described before between the normalized spectrogram of each rendition of the target syllable part and the baseline reference spectrogram.

We carried out a full clustering of all song syllables recorded from one day per week per bird, so we analysed 7 days per bird at maximum, *i.e*. one baseline day and 6 days post-deafening. To do so, song syllables were split based on threshold crossing on the root mean square (RMS) sound waveform, where the threshold was adjusted from one bird to another but kept constant for a given bird for all days analysed. Individual song syllables were manually sorted in different clusters according to their spectrographic similarity and position within the song motif. We then randomly extracted a sample of 200 renditions (maximum) of each syllable per week of experiment and per bird. From each song syllable rendition, including the entire target syllable, we computed the spectrogram divergence score as described above, but on the entire syllable (e.g. not as previously on a window of 48 ms), using our custom-made algorithms. We also used the Sound Analysis Pro program (Tchernichovski et al., 2000) to extract syllable duration, mean entropy and entropy variance. Mean entropy and entropy variance were computed as mean over the entire syllable. Note that because of a loss of data (hard drive failure) of the hearing birds, we were only able to compute the spectrogram divergence scores of the 48-ms target syllable and their number for this group of birds. For graphical representations at the population level, data of each feature were normalized using z-score values z:

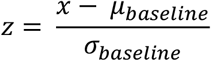

Where *x* is the value of the feature of interest for a specific syllable rendition, *µ* and *σ* are the mean and the standard deviation values, respectively, of the same feature computed on all syllable renditions on the baseline day.

### 5. Statistics

To evaluate whether the visual substitution task allowed deafened birds to maintain the target song syllable, we first computed the spectrogram divergence score of the 48-ms subpart of the target syllable and applied a linear mixed-effect model (LME) defined as:

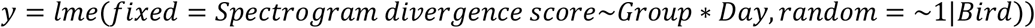

Where the factors *Group* and *Day* respectively accounts for the experimental groups (deaf LO, deaf no LO, deaf random LO and hearing birds) and days of experiment (from -5 to 35 days relative to the onset of the LO protocol).

In deafened birds, to determine whether the task impacted other acoustic features and other song syllables, for one day per week of experiment per bird, we extracted 200 renditions of each song syllable. For each acoustic feature (spectrogram divergence score, duration, mean entropy and entropy variance), we computed linear mixed-effect models on the raw measurements defined as:

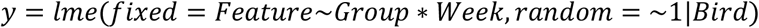

Where *Feature* corresponds to the one of the acoustic features measured, and the factors *Group* and *Week* respectively accounts for the experimental groups (deaf LO, deaf no LO and deaf random LO) and weeks of experiment (from -1 for pre-deafening, to 6 weeks post onset of the task).

For each LME, we computed an ANOVA on the output of the model followed, when appropriate, by post-hoc Tukey tests between significant factors.

All statistical tests were done using custom-written scripts in Matlab (v2023b) or R v4.4.1.

## Results

### 1. The spectrogram divergence score: a new measure of song stability

To examine whether deaf zebra finches are able to exploit a visual signal to compensate the post-deafening song degradation, we used a method based on the comparison of spectrograms, the spectrogram of the currently produced target syllable and a ‘reference spectrogram’ of this syllable. The reliable reference spectrogram was computed from all syllable renditions (>1000) of the syllable produced on a baseline day. At first, we ensured that this method allowed us to evaluate to what extent a syllable structure can be altered. To this end, we created samples of birdsongs that included various amount of acoustic distortion. They were built from songs of 19 hearing adult male zebra finches. We randomly extracted 200 renditions of an individual’s song motif produced in a single day and added a white noise (WN) of an intensity between 0 to 100% (step: 10%) such that the maximum sound amplitude never exceeds the one in the original song (Figure 1B). Spectrogram divergence scores were calculated from the comparison between the reference syllable spectrogram and the noisy syllable spectrograms. Score values appear to depend on the percentage of white noise intensity, increasing as intensity increases up to 40%, with little overlap between distributions of scores obtained for a given percentage and the slightly higher percentage, e.g. between 10% and 20% of WN in the signal. Beyond 50% of WN intensity, score values reached a plateau at a value of ∼0.2 (Fig 1Ci-Di). We also performed comparisons from entire song motifs, e.g. song motif 1 with 0% of WN *vs*. song motif 1 with 100% of WN, to measure the impact of white noise overlay. Spectrogram divergence scores for the entire motif also asymptotically increased as the WN intensity increased (Figure 1Cii-Dii). Importantly, they were similar to score values based on the analysis of the target syllable structure indicating that the degree of alteration of a selected target syllable may provide information of the distortion at the level of the entire song. Other possible measures of the degree of acoustic similarity between song motifs were the dissimilarity (see Methods for details) and the entropy scores provided by Sound Analysis Pro (SAP, Tchernichovski et al., 2000). The entropy score provides a measure of the randomness of the difference between acoustic signals. Computing the two scores from the samples of song motifs also resulted in quite similar positive and horizontal asymptotic curves (Figure 1 Ciii-Diii and Civ-Div). We can however note the great variability in SAP dissimilarity scores obtained by song motifs masked by WN at a given intensity, especially when the WN percentage is 20 or 30% and, consequently, the overlap between score distributions of song motifs masked by different intensities of WN. Also, the percentage of errors of computation using SAP increased with the level of WN added to the song motif while it remains zero using the spectrogram divergence scores (bar plots in green in Figure 1 Ciii-Diii). In spite of these differences, analyses based on spectrogram divergence score or SAP measures provided a similar picture of changes in song structure due to the addition of various levels of WN. This led us to use the spectrogram divergence score to measure the degree of similarity between syllables or songs motifs and to evaluate the degree of deafening-induced song degradation. Moreover, the spectrogram divergence score relies on a very rapid computation (<4ms) allowing us to deliver very rapidly a transient light-off extinction in an online syllable-contingent behavioural training paradigm.

**Figure 1:**
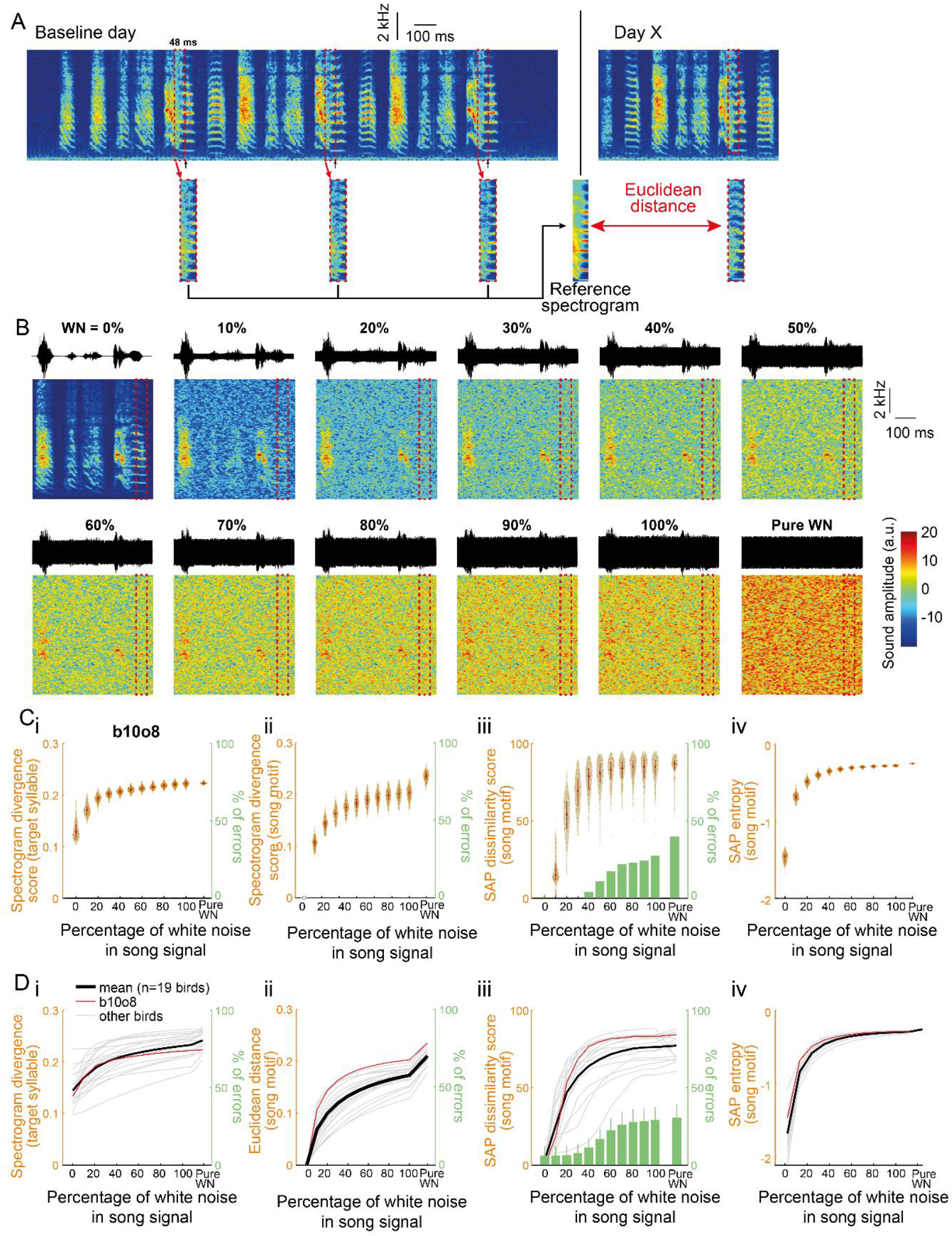
A new feature to assess sound degradation. **A**. Spectrogram of three consecutive motifs produced by a single bird on a baseline day. An algorithm is trained to detect a song syllable (black arrows) and to extract its spectrogram from the 48 ms window before the detection point (dashed red box). The median of all 48-ms syllable excerpts provided the “Reference spectrogram”. On the following days, the Euclidean distance was computed between the spectrograms of the targeted syllable and the Reference spectrogram to obtain the Spectrogram divergence score. **B**. Waveforms (top) and spectrograms (bottom) of the same song motif from a single bird (labeled b10o8) with increasing levels of white noise (WN) added to the signal and of pure white noise. The dashed red box highlights this bird’s 48-ms syllable excerpt used to assess the spectrogram divergence score. **Ci**. Violin plot showing the spectrogram divergence score (left y axis) computed between 200 randomly selected renditions of the 48-ms syllable excerpt for bird b10o8 with increasing levels of white noise in the signal (see B for spectrogram representation) and its reference spectrogram computed in the absence of white noise. White dots on the violin plots correspond to the mean spectrogram divergence scores. Note that there are no errors in the computation of the spectrogram divergence score (right y axis). **ii**. Euclidean distances between 200 randomly selected song motifs with the same motifs that include increasing levels of white noise in the signal or with pure white noise (left y axis). There is again no error in the computation of the score (right y axis). **iii**. Violin plots showing the dissimilarity score computed using Sound Analysis Pro (SAP; formula: 100-similarity score) between 200 randomly selected song motifs with the same motifs that include increasing levels of white noise in the signal or with pure white noise (left y axis). The percentage of errors of computation (when SAP failed in computing the similarity score between two songs) increases with the level of white noise added to the signal (green bar plot, right y axis). **iv**. Violin plot of the entropy between 200 randomly selected song motifs with the same motifs that include increasing levels of white noise in the signal or with pure white noise. **D**. Spectrogram divergence scores (**i**), Euclidean distance (**ii**), SAP dissimilarity score (**iii**) and SAP entropy (**iv**) computed as in C for 19 birds. Mean is shown with a thick black line and individual traces are in grey. Example bird from A, B, and C (b10o8) is shown in red.

### 2. Impact of the light-off exposure on the spectrogram divergence score in deafened birds

A previous experiment (Zai et al., 2020) revealed that deaf birds can exploit a visual signal to adapt the pitch of a song syllable. Here, we wondered whether deaf birds could take one step further in using this visual signal to offset the deafening-induced degradation of a given song syllable. We recorded the songs of 40 birds over a minimum of 7 consecutive weeks. A total of 33 birds were deafened after about one week of baseline song recording (Fig. 2A). Birds were trained according to an operant conditioning paradigm based on a computer program that detects a specific target song syllable, usually a harmonic-like syllable (Canopoli et al., 2014; Zai et al., 2020). Deafened birds were separated in three independent groups: deaf LO (n=13), deaf no LO (n=15) and deaf random LO (n=5). Deaf LO birds were trained by briefly switching off the light in the sound-isolation chamber whenever the spectrogram divergence score of the targeted syllable was below a threshold. The threshold for triggering the light extinction was adjusted on a daily basis to the median of the previous day. Deaf random LO birds were exposed to a similar amount of transient light extinction than deaf LO birds, but the visual signal did not depend on the syllable spectrogram divergence score. Deaf no LO and hearing birds were not exposed to transient light extinction during the entire course of the experiment.

**Figure 2:**
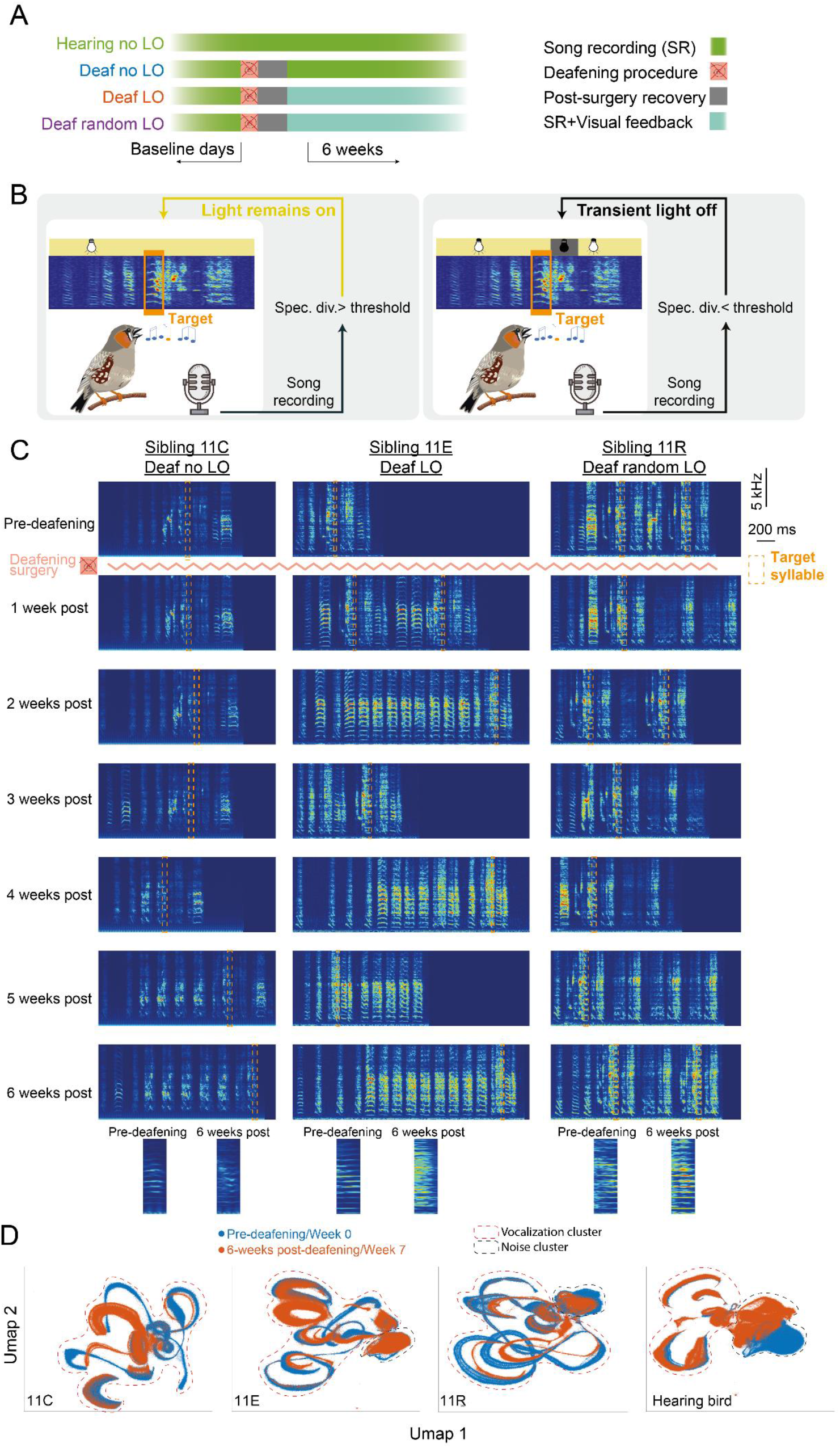
A behavioral task to counteract deafening induced song degradation. **A**. Birds were separated in four groups. All birds were isolated in a sound-proof chamber and their song was recorded on a few consecutive baseline days. Except the hearing birds (n=**7** birds), all the other birds were deafened following the procedure described in (Zai et al., 2020). After the deafening procedure, birds recovered for a few days before the onset of the behavioral paradigm. **B**. During the task, songs were recorded online and a target syllable specific for each bird was automatically detected. For deaf LO birds (n=13 birds), whenever the spectrogram divergence score was lower than a certain threshold, the housing light in the sound-proof chamber was transiently switched off for 200 ms. The light remained on otherwise. For deaf no LO birds (n=15 birds), the light was never switched off while for deaf random LO birds (n=5 birds), the light was transiently but randomly switched off once the bird produced the target syllable. **C**. Spectrograms of song motifs produced by three birds from the same clutch before and 1 to 6 weeks post-deafening and onset of the LO protocol. The target syllable for each bird is shown with a dashed orange box and zoomed in versions produced pre-deafening and 6-weeks post-deafening are highlighted. **D**. UMAP projections of all sounds recorded (i.e. birds vocalizations and cage noise) in the sound proof chamber for each example deafened bird and for a hearing control bird before deafening (blue) and 6 weeks post-deafening and onset of LO protocol for deafened birds, or 7 weeks later for the hearing bird. Clusters that include bird vocalizations (songs and calls) and noise are surrounded by dashed red and blue lines, respectively.

At first, we visually inspected spectrograms from songs recorded before and after deafening. As previously described, we observed a loss of song motif stereotypy occurring progressively over the six consecutive weeks of song recording in deaf birds. Fig. 2 shows spectrograms of song examples from three deaf birds, one per experimental group, before deafening and during the six weeks of the behavioural task. Importantly, these three example birds (a deaf no LO bird: 11C, a deaf LO bird: 11E and a deaf random LO bird: 11 R) belonged to the same clutch (clutch 11), so they were exposed to the same father’s song as juveniles. Deafening these birds resulted in a reduction in the degree of similarity between their songs, as indicated by the SAP similarity score computed from a subset of 200 songs recorded before and six weeks after the deafening procedure (see table 1 for details). We performed a UMAP analysis on the song recording datasets, selecting one day of the pre-deafening baseline recording period and one day of the sixth post-deafening week. UMAP graphs offer the advantage to represent many-dimensional data in just a few dimensions, e.g. two in Fig. 2D. Qualitatively, most of the clusters from songs produced by the birds before and six weeks after the deafening procedure show a low degree of overlap, so reflecting the overall modifications of the individual vocal repertoire. In comparison, the UMAP analysis performed in an example hearing bird shows that the vocal repertoire remained more stable than in deafened birds over a similar period of time (Fig. 2D).

**Table 1:** SAP similarity score between 3 siblings from the same clutch pre-deafening *vs* 6-weeks post-deafening.

| Birds | SAP similarity score (mean %+/-SD) pre-deafening | SAP similarity score (mean %+/-SD) 6-weeks post-deafening |
| --- | --- | --- |
| 11C (deaf no LO) vs 11E (deaf LO) | 33.2+/-11.1 | 29.3+/-6.9 |
| 11C vs 11R (deaf random LO) | 66.3+/-18.7 | 42.0+/-9.8 |
| 11R vs 11E | 60.9+/-9.6 | 48.5+/-9.8 |

Although visually informative, the UMAP analysis told nothing about changes in song structure. We first assessed whether the exposure to the behavioural training conditions differentially impacted the spectrogram divergence scores of the sibling of three birds (Fig 3A). The spectrogram divergence score increased over the days for the example Deaf no LO bird (11C bird : +23% between the baseline period and days 25 to 30 post start of the LO protocol) while it remains rather stable for the two examples of deaf LO (11E bird : +14.5%) and deaf random LO (11R bird: +12.3%) birds, even though the number of renditions of the target syllable sang by the example deaf LO bird decreased more rapidly than for his control sibling. At the population level, the spectrogram divergence scores of the 48-ms subpart of the target syllable remained stable for hearing birds while it regularly increased for deafened birds of the three groups (Fig. 3B). Note that even if the LO protocol lasted 6 consecutive weeks, we restricted the analysis at the population level to 4 consecutive weeks in order to include all the birds in the analysis. Indeed, over time, the target syllable was less and less detected in deafened birds. The difference between hearing and deafened birds increased across days revealing a cumulative effect and thus a gradual degradation of the overall accuracy of the 48-ms target song syllable in all deafened birds. We computed the spectrogram divergence scores on all detected renditions of the 48-ms target syllable and normalized (Z-score) the data to the last day before the onset of the LO protocol. A linear mixed effect model (LME) on the spectrogram divergence scores calculated over 35 consecutive days, including 5 days of baseline, with the birds as a nesting factor, revealed significant changes over days (F_34,1066_=22.18, *p*<0.001) and significant differences between groups (F_3,36_=3.30, *p*<0.04),. A post-hoc analysis (Tukey) on the group effect revealed greater changes in the syllable structure in deafened birds which were not exposed to the LO procedure compared to hearing birds (Deaf no LO vs. hearing birds; t-test_36_=3.08, *p*<0.02), and compared to deafened birds exposed to the LO procedure (Deaf LO vs. Deaf no-LO birds; t-test_36_= 2.75, *p*<0.05). Comparisons based on scores computed on the target syllable produced by birds of the random LO group, which included only five birds, did not reveal any difference with the other groups. These results thus highlight a partial maintenance, or at least a slower deafening-related degradation of the 48-ms target syllables due to the exposure to the visual signal.

**Figure 3:**
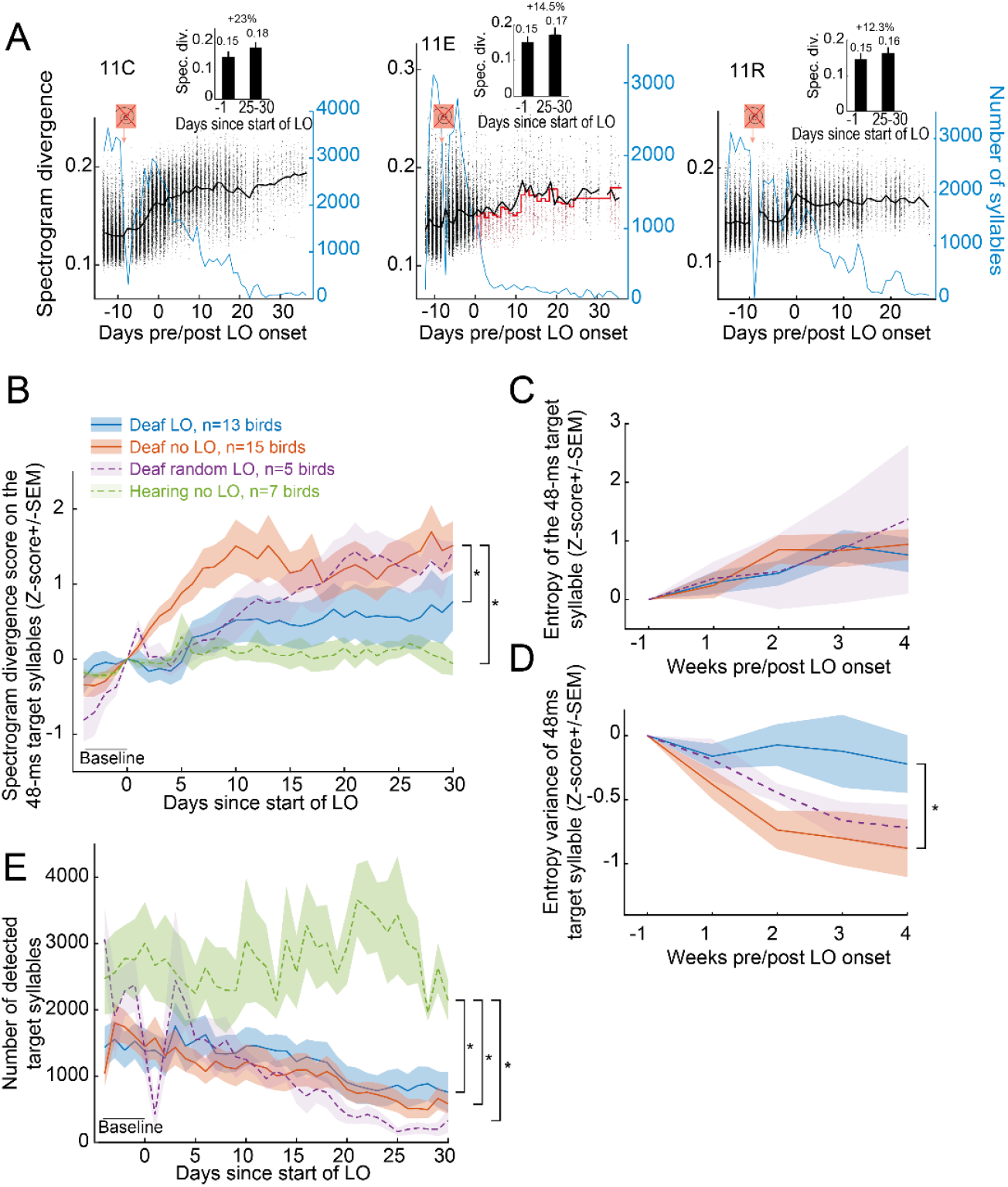
Transient light extinction contingent on the spectrogram divergence score slows syllable degradation in deaf birds. **A**. Variations in spectrogram divergence score (black dots, left y axis) and the number of detected target syllable (blue line, right y axis) over days for three example deafened birds (same birds as in Fig. 2): the deafened 11C no LO (left), the deafened 11E LO (middle) and the deafened 11R random LO bird (right). Black line indicates the mean value of the spectrogram divergence score with the dots representing the raw values calculated for each detected target syllable. Red line for the deafened 11E LO bird indicates the threshold value used for triggering the transient light extinction. The symbol in orange indicates when deafening procedure occurred. The LO exposure started the day zero, with the same day used for both the 11C and the 11E birds. Inserts show the mean (+/-SEM) spectrogram divergence score over the last day before the onset of the LO protocol and days 25-30 after the onset of the LO protocol. Above the percent of change. **B**. Normalized spectrogram divergence score (mean Z-score+/-SEM) computed on all detected renditions of the 48-ms target syllable for all birds before and during the LO protocol. **C-D**. Entropy (C) and entropy variance (D) computed on a subset of 200 randomly selected renditions per week per bird of the 48-ms target syllable. **E**. The number of target syllables automatically detected decreased for all deafened birds. *, significant group effect, p<0.05 (see text for details).

Entropy and entropy variance were previously used to evaluate post-deafening song degradation (Horita et al., 2008). In order to assess whether the effect of the LO exposure on syllable structure in deaf birds was also observed using the entropy and the entropy variance as measures,, we randomly extracted 200 renditions of the target syllable (48 ms) per week and per deaf bird (Fig 3C-D). A LME computed on these two measures revealed significant changes of both –entropy and entropy variance over weeks (entropy: F_4,115_=9.99, *p<*0.001; entropy variance: F_4,115_=10.83, *p<*0.001). The target syllable entropy did not depend on the use or not of light extinction (no group effect: F_2,30_=0.09, *p*=0.91; no group*week interaction : F_8,115_=0.51, *p*=0.85). In contrast, the entropy variance was affected by the behavioural protocol (group*week interaction: F_8,115_=2.10, *p*=0.04), with a significant difference between birds contingently exposed to the light extinction compared to those not exposed to the visual signal (Deaf LO *vs*. Deaf no LO; t-test_36_= 2.64, *p*=0.03) providing an additional evidence that exposure to the visual signal affected the deafened-induced degradation of the target syllable, at least the 48 ms period used as reference to deliver the visual signal. The entropy variance of the syllable target of the five birds of the group randomly exposed to the LO procedure did not differ from that of the two other groups of deaf birds.

In a previous study, the exposure to a pitch-contingent LO protocol was found to affect the singing activity of deaf birds (Zai et al., 2020). The pitch-contingent delivery of a transient light extinction led deafened birds to show a high motivation to sing (Zai et al., 2020). We examined here whether the singing rate varied depending on the contingency of the LO protocol. We estimated the number of motifs sung by individuals on a daily basis from the number of renditions of the target syllable, considering that the birds always included the target song syllable within each song motif. As shown by the three examples of deaf birds (blue line in Fig. 3A), birds sang less and less over the six weeks of song recording (Fig. 3E). A LME computed on the number of target syllables automatically detected, with the birds as a nesting factor, revealed significant group effect (F_3,36_=5.29, *p*<0.005), day effect (F_34,1066_=12.80, *p*<0.001) and day*group interaction (F_102,1066_=1.77, *p*<0.001). The number of detected target syllables showed a gradual decrease over days in the three groups of deafened birds but not in the group of hearing birds (post-hoc Tukey test; Deaf LO *vs*. Hearing birds: t-test_36_ =3.51, *p*<0.007; Deaf no LO *vs*. Hearing birds: t-test_36_ =4.09, *p*<0.002; Deaf random LO *vs*. Hearing birds: t-test_36_ =3.02, *p*<0.03) but with no significant differences between groups of the deafened birds. The singing activity showed thus a similar gradual decrease in the three groups of deafened birds, so independently of the contingency of the LO protocol with the spectrogram divergence score (Fig. 3E).

Computation of the spectrogram divergence score was computed on a window of 48 ms of the target syllable. The syllable duration was longer leaving unclear whether the slower degradation of the target syllable structure exhibited by deaf LO birds was limited to the 48ms window or extended to the entire target syllable, and beyond, to the entire song. We computed offline the spectrogram divergence scores from the entire target syllable and all other song syllables (“non-target syllables”). We established a reference syllable spectrogram for each syllable type found in the individual’s song motif, from 200 randomly selected renditions of each syllable during a baseline day. We also randomly selected 200 syllable renditions per week over the four weeks of the post-deafening period to compute spectrogram divergence scores. As shown by the Fig. 4A, spectrogram divergence scores of the entire target syllables as well as other syllables increased over weeks (week effect; target syllable: F_4,120_=77.94, *p<*0.001; non-target syllables: F_4,509_=120.70, *p<*0.001).

**Figure 4:**
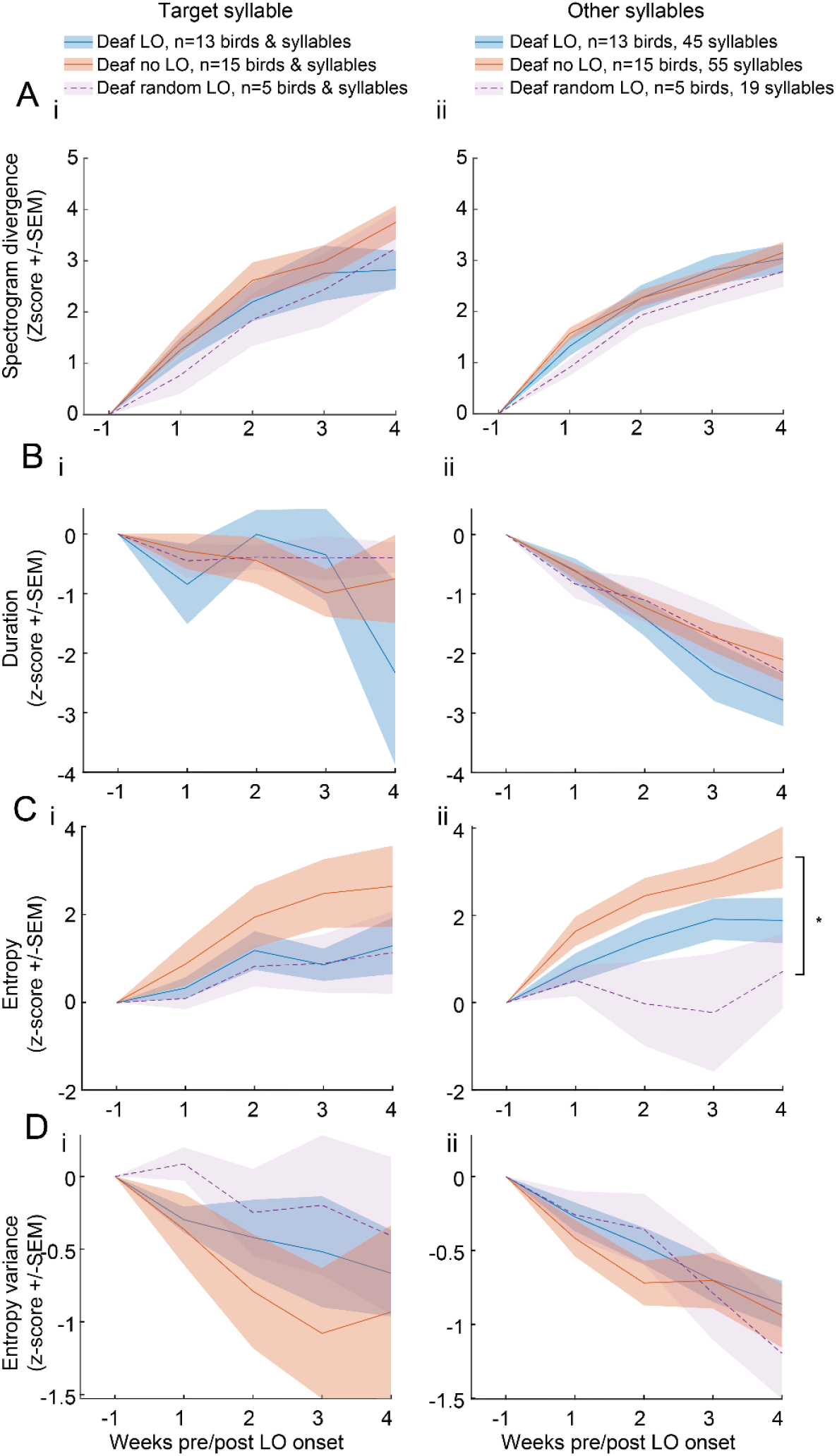
Various acoustic features show no clear evidence of an impact of a LO exposure on the structure of the entire target syllables or of other song syllables. Acoustic measures were computed on subsets of 200 randomly selected target (left panel) and non-target (right panel) syllables per bird and per week. Spectrogram divergence score (**A)**, syllable duration (**B**), entropy (**C**) and entropy variance (**D**) of the entire target (i) or non-target (ii) syllables over the 6 weeks after the onset of the LO protocol for deaf LO birds. Spectrogram divergence scores evolved within a similar range for the three groups of deafened birds. *, significant group effect, p<0.05 (see text for details).

When we compared the extent of changes in spectrogram divergence score values over time between the three groups of deafened birds, the target syllable as well the non-target syllables did not reveal any effect of the behavioural protocols (target syllable, group effect: F_2,30_=0.95, *p*=0.40; group * week interaction: F_8,120_=1.55, *p*=0.15; non-target syllable, group effect: F_2,29_=1.14, *p*=0.33; group * week interaction: F_8,509_=0.42, *p*=0.91). Spectrogram divergence scores therefore suggest that the impact of the exposure to the light off procedure did not extent the 48ms period used to compute the threshold value for triggering the visual signal.

We also analysed whether the duration, entropy, entropy variance of the syllables changed over weeks, depending on the behavioural protocol. We used the same dataset of 200 randomly selected renditions of each syllable type per week and per bird. The duration of the target syllable was rather stable over weeks, at least the first three weeks (Fig 4B; F_4,117_=2.01, *p<*0.10), while the duration of non-target syllables decreased (F_4,521_=37.83, *p<*0.001) with no difference in the course of changes between the three groups of deafened birds (target syllable: F_2,30_=0.11, *p*=0.89; non-target syllables: F_2,29_=0.62, *p*=0.54). The entropy of target and non-target syllables increased over weeks (target syllable: F_4,117_=15.10, *p<*0.001; non-target syllables: F_4,521_=11.70, *p<*0.001; Fig. 4C) while entropy variance regularly decreased (target syllable: F_4,117_=5.42, *p<*0.001; non-target syllables: F_4,521_=17.48, *p<*0.001; Fig. 4D) indicating that song syllables gradually became uniformly noisier. The course of changes in the entropy of the target syllable did not reveal any difference between the three groups of deafened birds (group effect: F_2,30_=1.21, *p*=0.31; group*week interaction: F_8,117_=1.82, *p*=0.32). In contrast, the course of the entropy of other song syllables varied depending on the behavioural protocol used (group effect: F_2,29_=4.18, *p*<0.03). The entropy differed significantly between deaf no LO and deaf random LO birds (post-hoc Tukey test, deaf no LO vs. random LO birds: t_29_=2.83, *p*<0.03), with no difference between deaf LO birds and the two other groups.

Only the entropy revealed difference depending on the behavioural protocol used. The entropy variance of both the entire target syllable and the set of other syllables decreased over weeks, with no difference in the time course between groups (target syllable; group effect : F_2,30_=0.63, *p*=0.54; group*week interaction: F_8,117_=0.45, *p*=0.88; other syllables; group effect: F_2,29_=0.18, *p=*0.83; group*week interaction: F_8,521_=0.49, *p*=0.86).

## Discussion

In both humans and songbirds, loss of auditory feedback in adulthood leads to progressive loss of precise vocal control (Waldstein, 1990; Lane and Webster, 1991; Nordeen and Nordeen, 1992; Lombardino and Nottebohm, 2000; Horita et al., 2008; Tschida and Mooney, 2012). This observation led to the hypothesis that auditory feedback is necessary for the maintenance of speech production. However, previous studies showed that both deafened and hearing birds can adjust the pitch of a selected song syllable using non-auditory feedback signals, including a light extinction contingent to the pitch of the target syllable (Zai et al., 2020; McGregor et al., 2022). Here, we provide evidence that deafened birds can also use light-off signal to slowdown deafness-induced song degradation of a song syllable. A deterioration in the song structure was exhibited by all deafened birds. The deterioration of the structure of a given syllable, measured thanks to the spectrogram divergence score, however, appeared to be reduced in deafened birds exposed to the light off procedure in comparison to deafened birds that did not experienced visual signals. The present study therefore suggest that visual information could, to some extent, be used by deafened birds to control the structure of a given song syllable

Previous studies reported that deafened birds are able to control the pitch of a selected syllable in a behavioral training paradigm that relies on the contingent delivery of a sensory signal depending on the pitch value of the syllable (refs). This control raised the question to know whether a similar training paradigm could allow birds to counteract consequences of the loss of auditory signals and slowdown the deafening-induced degradation of a syllable. The aim of the present study was to address this issue. To our knowledge, compensating for deafening-induced song degradation after deafness had never been attempted. However, deafening-induced song degradation includes a wide range of spectro-temporal changes in the structure of song syllables (Nordeen and Nordeen, 1992; Wang et al., 1999; Brainard and Doupe, 2001; Horita et al., 2008). Also, prior the present study, no acoustic measurement of the entire syllable structure had been carried out so quickly that it could be used in a behavioural task. In previous studies, song syllable degradation was quantified either visually by the observation of spectrograms, or by the quantification of some acoustic features, including mean entropy and entropy variance that provide good estimates (Horita et al., 2008; Pytte et al., 2012; Tschida and Mooney, 2012). These measures were, however, performed offline. Therefore, training birds in a behavioral paradigm to enable them to maintain potentially the overall structure of one of their song syllables by specifically targeting it, required to be innovative. In order to quantify deafness-induced song degradation more globally, *i.e* on the basis of multiple acoustic features, and very quickly, we developed a new method based on the computation of the spectrogram divergence score. This measure can be seen as rapidly reflecting the overall stability of a song syllable and post-deafening song changes. The spectrogram divergence score values remained stable in hearing birds over weeks. Deafening induced striking changes in song and syllable structure and song spectrograms became gradually less similar to the song produced before. Consistently, spectrogram divergence scores in deafening birds that were not exposed to the light off extinction protocol gradually increased over weeks, in a range of values that significantly differed from score values computed from spectrograms of hearing birds. Also, this measure allowed us to capture fine changes in syllable structure since spectrogram divergence score values revealed a deafening-induced impact from the first weeks. The spectrogram divergence score that has the advantage of analyzing the global structure of vocal signals in a way that is fast enough to be used online for a behavioral task, therefore proved to be a reliable and effective measure of song degradation. This method could considerably contribute to quantify short and long-term degradation of syllable structure in future studies and could therefore have relevance to the field of vocal plasticity.

On the basis of both the spectrogram divergence score and entropy variance values, deafened birds exposed to the transient light extinction contingently to the computation of the spectrogram divergence score retained to a certain extent the overall structure of the target syllable. Both the use of a visual signal and the contingency of its delivery might be crucial in enabling the birds to control, at least in part, the structure of the target syllable. Deafened birds that were not exposed to light extinction exhibited a faster and a more severe degradation of the target syllable structure. The amount of degradation of the target syllable appeared to be moderate in birds randomly exposed to the visual signal. It remains, however, to examine to what extent the contingency of the visual cue delivery guides the control of the target syllable structure. No clear difference in the severity of the degradation allows distinguishing birds contingently exposed to light extinction from birds exposed randomly. The present study, nevertheless, provides evidence of a weaker impact of the deafness when a visual cue was delivered. This suggests that deaf birds are able to exploit a transient light extinction signal not only to adjust the pitch of a target syllable (Zai et al., 2020), but also to maintain at least partially its spectro-temporal structure. Consequently, even though the auditory feedback is critical to provide evaluation of what has been vocally produced, information of another sensory modality can influence the song production. Yet, the impact reported here on syllable structure maintenance appears to not extent beyond the 48 ms period used to calculate the similarity score between the reference spectrogram and the spectrogram of the new rendition of the syllable target. No difference was found in divergence spectrogram scores computed from either the entire target syllable or other non-targeted syllables between deaf birds exposed or not to the light-extinction. Syllable duration also did not exhibit any difference. Entropy measures only indicate a possible effect of the behavioral conditions. These results are consistent with previous studies investigating the impact of a pitch-shifting paradigm on the acoustic structure of the syllable target (Tumer and Brainard, 2007; Canopoli et al., 2014; Pehlevan et al., 2018; Zai et al., 2020). Even if deaf birds are able to modify the fundamental frequency value of the target syllable in a time-period that extended beyond the time window used to calculate the pitch, changes only occur in a very narrow time-period around the pitch calculation time window (Canopoli et al., 2014; Zai et al., 2020). Other song syllables do not show any changes in their fundamental frequency or other spectro-temporal features (Tumer & Brainard, 2007). Additionally, the behavioral training task used in the present study, because it was based on the overall acoustic structure of the syllable and not on a single acoustic parameter, can be considered as probably more complex or difficult for the birds than the pitch-shifting behavioral task. The fundamental frequency may only be controlled by one set of muscles surrounding the syrinx (Larsen and Goller, 2002; Goller and Riede, 2013). To maintain all the acoustic parameters of a given syllable, it is likely that a greater number of muscles need to be controlled.

It remains now to investigate changes in neural networks and to identify specific neural plasticity mechanisms underlying the integration of visual signals used to replace auditory information in deaf birds. Given brain reorganizations that occur as a result of deafness (Pytte et al., 2010; Tschida and Mooney, 2012; Zhou et al., 2017), the behavioral training task we used here, could therefore have impacted, at least in part, these reorganizations. It would be interesting in order to study multisensory interactions that can take part in vocal production, to investigate how visual signals contingent on one acoustic feature, *e.g*. the fundamental frequency, of a song syllable affect brain circuitry. In 1985, a study showed that neurons in the sensorimotor HVC nucleus may exhibit changes in their activity in response to visual stimuli, suggesting that the song system receives visual information (Bischof and Engelage, 1985). We have shown previously that a lesion of the songbird basal ganglia (Area X) abolishes the ability of deafened birds to adjust their song syllable pitch using a transient light extinction (Zai et al., 2020). In order to exploit transient light stimulus, birds probably use information that enables them to control motor output so that they can deliberately modify it. Information may be proprioceptive, but could also be based on an internal motor copy of the song command from motor centers to Area X (i.e. efferent copy, Giret et al., 2014). This copy, in conjunction to the release of reinforcing signals (*e.g*. dopamine) into Area X during the contingent sensory feedback (Gadagkar et al., 2016; Roeser et al., 2023), could provide, in parallel, information about the motor command in sensory regions. A future challenging question is how visual information in deafened birds affects song system processing to guide the vocal motor control and counteract the deafening-induced syllable degradation in absence of auditory feedback (Rolland et al., 2022). Future works should investigate to what extent the vocal behavior of songbirds relies on visual information processing by song system nuclei.

## Acknowledgments

This work was supported by the *Centre National de la Recherche Scientifique* and the University of Paris Saclay. M.R. was supported by the *Ministère de la Recherche, de l’Enseignement supérieur et de l’Innovation*. We thank Sophie Cavé-Lopez for help in conducting the experiments. We thank Mélanie Dumont and Caroline Rousseau for taking care of the songbird facility.

## Notes

### Competing Interest Statement

The authors have declared no competing interest.

